# Splicing of ultraconserved poison exons controls mitotic fidelity and stem cell viability

**DOI:** 10.64898/2026.05.19.726407

**Authors:** Nathan K. Leclair, Mattia Brugiolo, Isha Walawalkar, Marina Yurieva, Hyeon Gu Kang, Ryan Englander, Mallory Ryan, Caleb Heffner, Justin A. McDonough, William C. Skarnes, Steve Murray, Olga Anczuków

## Abstract

SR proteins are essential splicing regulators whose expression is controlled in part through poison exons (PEs) — ultraconserved non-coding exons that trigger nonsense-mediated decay — yet the biological functions of these elements remain undefined. Here, we show that homozygous deletion of *SRSF3-PE* or *TRA2*β*-PE* is selected against in mouse embryos and human induced pluripotent stem cells (iPSCs), and that conditional PE deletion causes apoptotic death in iPSCs but is tolerated in post-mitotic neurons, revealing a proliferative-state-specific requirement. Mechanistically, PE deletion elevates SR protein levels, triggers widespread splicing dysregulation, and disrupts the correct splicing of a mitotic gene network associated with spindle defects and mitotic errors. These findings establish ultraconserved poison exons as essential regulators of mitotic splicing fidelity and stem cell viability.

## Introduction

The human genome harbors 481 ultraconserved elements (UCEs), DNA sequences of ≥200 base pairs that are 100% identical across human, mouse, and rat genomes (*1*). This degree of constraint is extraordinary: it exceeds the conservation of protein-coding sequences and implies that even a single nucleotide substitution over ∼80 million years of mammalian evolution has been selectively lethal. Yet despite two decades of study, no unifying biological function has been assigned to UCEs. The vast majority reside in non-coding regions, overlapping enhancers or gene bodies of RNA-processing factors, and targeted deletions of individual UCEs in mice produced no overt phenotype in early studies (*2–4*), deepening rather than resolving the mystery of their extreme conservation.

A subset of UCEs, however, occupies a functionally compelling position: they span or embed poison exons (PEs) within transcripts encoding serine-arginine rich (SR) splicing factors, a family of ubiquitously expressed RNA-binding proteins (RBPs) that act as master regulators of alternative splicing (AS) and indispensable for embryonic development, tissue homeostasis, and cancer cell survival (*5–13*). PEs are non-coding exons whose inclusion introduces a premature termination codon (PTC), triggering transcript degradation via nonsense-mediated decay (NMD). Through this coupled alternative splicing-NMD (AS-NMD) mechanism, SR proteins autoregulate their own expression and cross-regulate one another, maintaining precise splicing-factor stoichiometry across cell types and developmental states (*8, 9, 14–17*). Strikingly, the PE-containing UCEs in SR protein transcripts can span >500 base pairs and display higher nucleotide-level conservation than flanking coding exons — a phylogenetic signal suggesting that these regulatory sequences are under more intense selection than surrounding protein-coding sequences.

This study focuses on two proto-typical SR proteins, SRSF3 and TRA2β, both of which are ubiquitously expressed yet functionally non-redundant, with distinct and context-specific roles in development and disease (*18–22*). SRSF3 is required for cardiac and hepatic development, metabolic homeostasis, and pluripotency — including nuclear export of *NANOG* mRNA and AS regulation of pluripotency-associated genes (*18, 23–31*). TRA2β regulates somite and brain development, and neuronal function (*32–34*). Disruption of either protein has broad pathological consequences: reduced SRSF3 expression drives insulin resistance and liver disease, while TRA2β dysregulation is linked to neurodegeneration and obesity (*5, 26, 28, 35*). Additionally, both SRSF3 and TRA2β function as oncoproteins — elevated in lung, breast, ovarian, gastric, bladder, colon, hepatocellular, and head and neck cancers — where their overexpression promotes transformation and metastasis, and their depletion impairs proliferation (*6, 11, 36–40*). Recent work, including from our group, demonstrates that *SRSF3-PE* and *TRA2*β*-PE* inclusion is dynamically regulated during differentiation and disease including cancer, cardiac disorders, and immune activation, and that reduced PE inclusion correlates with worse patient outcomes in multiple cancer types — establishing *PE*-mediated AS-NMD as a functionally relevant regulatory axis (*7–9, 13, 17, 24, 35*). However, despite drastic phenotypes in tissue maintenance and disease states when PE splicing becomes disrupted, as well as contributions of their parent SR protein to differentiation and development, little is known about how these PEs contribute to stem cell physiology and early organism development.

Here, we demonstrate that homozygote deletions of *SRSF3-PE* and *TRA2*β*-PE* are selected against in mouse zygotes and human iPSCs, and that PEs are required for iPSC viability but dispensable in post-mitotic neurons. PE deletion triggers transcriptomic and splicing dysregulation, disrupts SF cross-regulatory networks, and impairs splicing of mitosis-associated genes, culminating in mitotic failure in proliferating cells, revealing ultraconserved poison exons as essential guardians of stem cell identity and developmental integrity.

## Results

### Homozygous deletion of ultraconserved poison exons are selected against in mouse embryos and human iPSCs

To investigate the role of ultraconserved PEs in mammalian development and pluripotent cells, we first chose to model PE loss through genomic deletion of the PE and flanking ultraconserved intronic sequences, without broader gene disruption. This approach is mechanistically distinct from full gene knockout models of *Srsf3* or *Tra2*β, which abolish protein-coding transcript expression and SR protein production and are associated with embryonic lethality in mice (*19, 22*). PE-specific deletion has the opposite molecular consequence: by removing the non-coding PE-containing transcript while leaving the protein-coding isoform intact, it abrogates AS-NMD-mediated degradation and is predicted to increase, rather than decrease, SR protein levels. We therefore used CRISPR/Cas9-mediated PE deletion in two independent model systems to test its impact on normal development and stem cell viability.

First, we electroporated paired-guide RNAs (pgRNAs) flanking the ultraconserved *Srsf3* or *Tra2*β regions along with Cas9 protein into mouse zygotes, which were cultured to the blastocyst stage or immediately transferred into donor mice for gestation to embryonic day 18.5 (E18.5), using protocols optimized by the Knockout Mouse Phenotyping Project and the International Mouse Phenotyping Consortium which routinely yield high biallelic rates at neutral loci (*41–44*) (**Fig. S1A**). Next, PE deletion was quantified by droplet digital PCR across blastocysts and day 18.5 (E18.5) embryos (**Fig.S1B**). Of the 101 blastocysts and embryos genotyped for *Srsf3*-PE and 82 for *Tra2*β-PE, only 3 and 2, respectively, harbored full homozygous deletions, far below the ∼50% biallelic deletion efficiency expected for non-essential loci using this approach (*45*). The predominance of heterozygous and mosaic embryos suggests that homozygous loss of these ultraconserved regions is under strong negative selection during early mouse embryogenesis.

To test whether this essentiality extends to a human model system, we delivered Cas9 mRNA along with pgRNAs targeting the *SRSF3-PE* or *TRA2*β*-PE* and ultraconserved regions into KOLF2.1J human iPSCs (**Fig.1A-B**). In parallel, to isolate the effect of PE skipping from that of a deletion at the genomic locus, we introduced point mutations inactivating the 3’ splice site (3’SS) of each PE (**Fig.1B**). Among 139 *SRSF3* and 144 *TRA2*β iPSC clones screened for genomic PE deletions, no viable homozygous clones were recovered (**Fig.1C**). For the 3’SS point mutations, no viable homozygous clones were recovered for *TRA2*β*-PE-3’SS^mut/WT^* but we obtained 19 viable homozygous *SRSF3-PE-3’SS^mut/mut^*clones (**Fig.1C**). Interestingly, RT-PCR of *SRSF3-PE* splicing in these clones revealed a smaller band and Sanger sequencing confirmed utilization of a cryptic alternative 3’SS located 43 bp downstream of the mutated canonical site, re-creating a structurally distinct, but functionally identical PE (**Fig.1C**; **Fig.S1C-D**). This spontaneous 3’SS rescue indicates that iPSCs are not only unable to tolerate PE loss, but actively select against it at the sequence level, preserving the PE inclusion mechanism even when the canonical splice site is destroyed. Together, these data establish that biallelic loss of *SRSF3*-PE or *TRA2*β-PE is strongly disadvantaged in both mouse embryos and human iPSCs, implying an essential function at the earliest stages of mammalian development.

**Figure 1:**
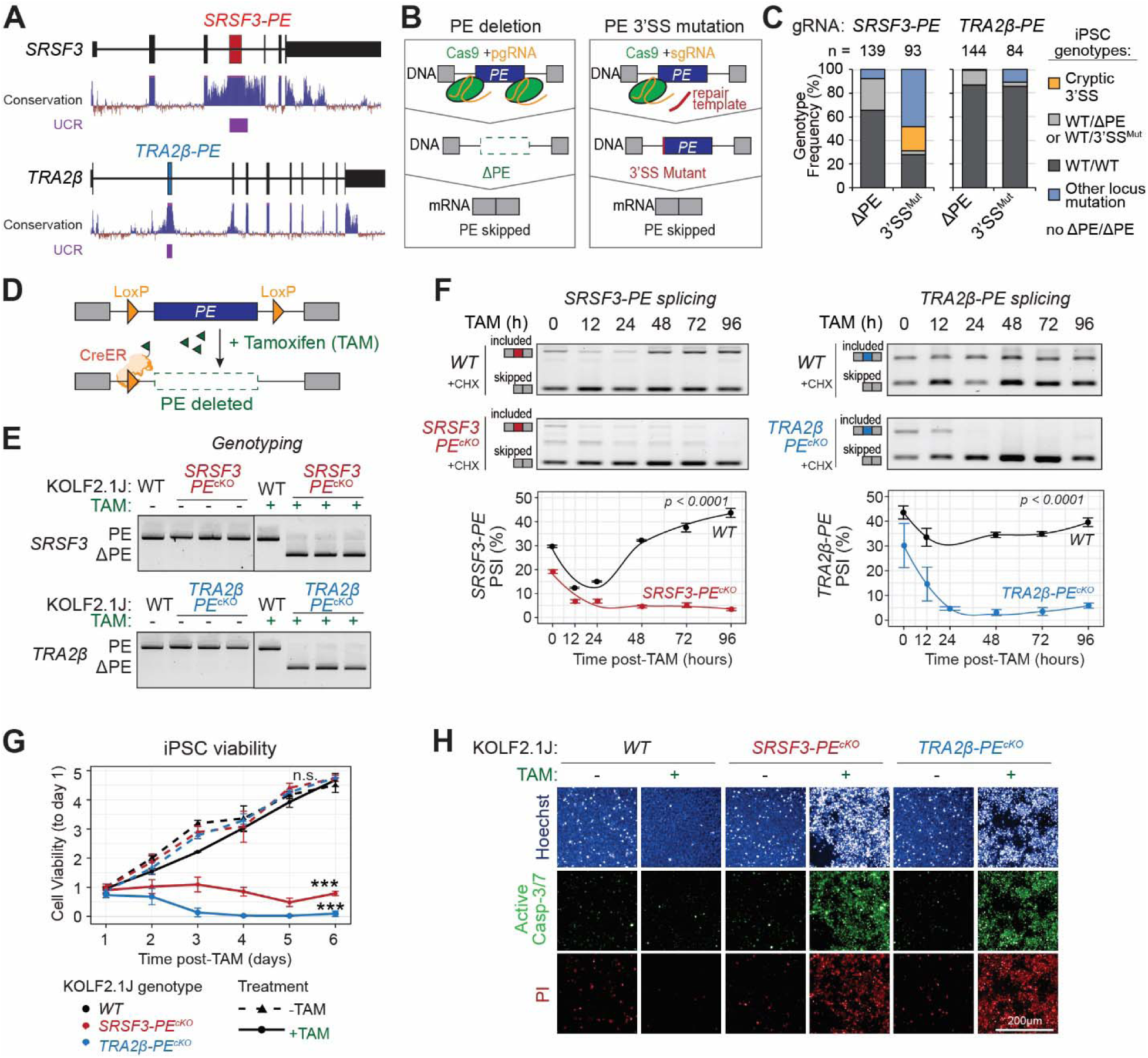
*SRSF3-PE* and *TRA2*β*-PE* are essential for human iPSC viability. **(A)** Human *SRSF3* and *TRA2*β gene structures showing conservation tracks and highlighting ultraconserved regions (UCR). **(B,C)** Schematic of the CRISPR/Cas9 approach to delete *SRSF3*-*PE* or *TRA2*β-*PE* using paired gRNAs (pgRNAs) (left), or to mutate and abolish the *PE* 3’SS using a single gRNA with a repair template (right) (**B**). Genotype frequency of isolated viable KOLF2.1J human iPSCs clones with stable *PE* deletion or 3’SS mutations (**C**). Total number of viable genotyped clones is indicated on top. **(D,E)** Conditional *PE* knockout model (*PE^cKO^*) in CreERT2 human iPSCs, in which addition of tamoxifen (TAM) triggers nuclear CreERT2 localization, recombination of LoxP sites flanking the *PE* sequences of *SRSF3* or *TRA2*β, and homozygous *PE* deletion (**D**). Genotyping of *SRSF3* or *TRA2*β loci in wild-type (WT), *SRSF3-PE^cKO^*, and *TRA2*β*-PE^cKO^* KOLF2.1J-CreERT2 iPSCs treated with (+) or without (-) TAM using primers located on each side of the PE (n=3 independent clones) (**E**). **(F)** *PE* splicing in WT, *SRSF3-PE^cKO^*, and *TRA2*β*-PE^cKO^* KOLF2.1J-CreERT2 iPSCs treated with TAM for indicated time points is measured using semi-quantitative RT-PCR using primers that amplify the included and skipped isoforms, quantified as percent spliced-in (PSI). Cells are treated with cycloheximide (+CHX) for 6h prior to collection to stabilize NMD-sensitive PE-containing transcripts. Representative images are shown along with quantification (n=3 independent clones, mean±SD; two-way ANNOVA with Tukey’s post-hoc adjustment). **(G)** Cell viability of WT, *SRSF3-PE^cKO^*, and *TRA2*β*-PE^cKO^* KOLF2.1J-CreERT2 iPSCs with (+) or without (-) TAM for indicated time points is assessed using cell titer glow assay and normalized to day 1 (n=3 independent clones, mean±SD; two-way ANNOVA with Tukey’s post-hoc adjustment; *P*-values untreated to TAM by genotype). **(H)** Cell death in *WT, SRSF3-PE^cKO^*, and *TRA2*β*-PE^cKO^* KOLF2.1J-CreERT2 iPSCs with (+) or without (-) TAM for 96h is assessed by measuring caspase-3/7 positive cells and stained with propidium iodide (PI). Nuclei are co-stained with Hoechst. Scale bar: 200µm.

### Conditional deletion of *SRSF3-PE* or *TRA2*β*-PE* is lethal in human iPSCs

Because homozygous PE deletions were not recovered by standard CRISPR editing, we developed iPSC lines with an inducible Cre/loxP conditional knockout (cKO) system to achieve controlled, temporally defined PE deletion. LoxP sites were inserted flanking the ultraconserved regions of *SRSF3* or *TRA2*β (encompassing the PE and ∼100-200 bp of conserved flanking intronic sequence) in KOLF2.1J iPSCs stably expressing a tamoxifen (TAM)-regulated Cre recombinase-estrogen receptor fusion (CreERT2) from a safe harbor locus (*46*), generating *SRSF3-PE^cKO^* and *TRA2*β*-PE^cKO^* lines. At baseline, CreERT2 is cytoplasmic and inactive; TAM addition drives nuclear translocation and loxP-mediated excision of the PEs (**Fig.1D-E**).

TAM treatment produced rapid, complete PE deletion, leading to loss of both PE allele and full loss of PE inclusion at the RNA level within 24 hours in three independent *SRSF3-PE^cKO^* or *TRA2*β*-PE^cKO^* clones, with no effect in wild-type (WT) controls (**Fig.1E-F**). PE deletion was accompanied by a 2- to 3.5-fold increase in SRSF3 and TRA2β protein levels within 48 hours (**Fig.2B**), consistent with loss of AS-NMD-mediated autoregulation. Critically, TAM-treated *SRSF3-PE^cKO^* or *TRA2*β*-PE^cKO^* iPSCs underwent progressive cell death within 3-6 days; viable cultures with stable biallelic PE deletions could not be maintained beyond this window (**Fig.1G**). Cell death was attributable to apoptosis, as evidenced by caspase-3/7 activation and propidium iodide stain in PE-deleted but not WT TAM-treated iPSCs (**Fig.1H**). These phenotypes were independently replicated in the WTC11 iPSC line (**Fig.S2**), confirming that PE essentiality is not a clone- or line-specific artifact but likely a general property of human pluripotent cells.

**Figure 2:**
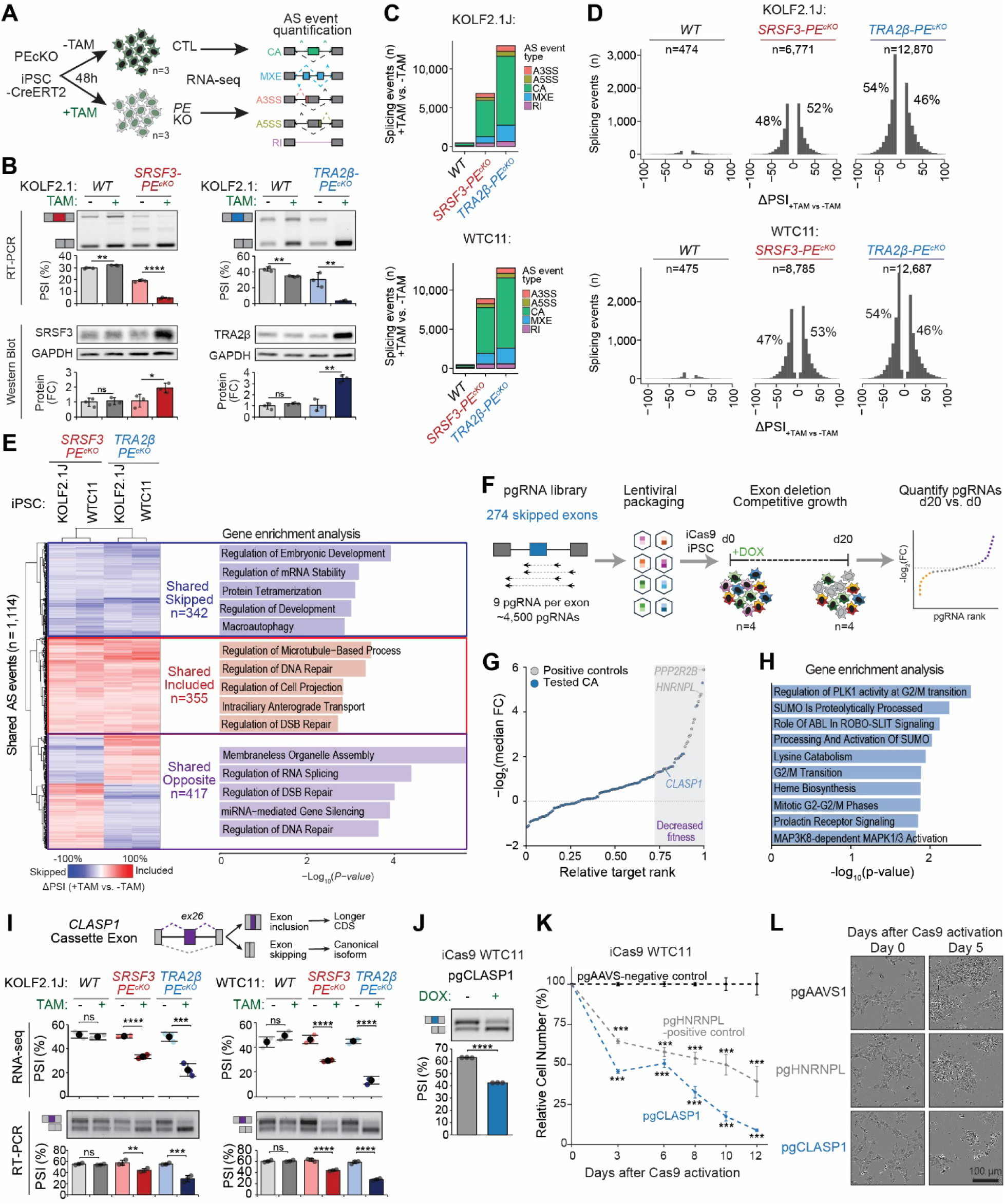
*SRSF3-PE* and *TRA2*β*-PE* deletions induce changes in iPSC splicing programs. **(A)** RNA-sequencing is performed on *WT*, *SRSF3-PE^cKO^*, and *TRA2*β*-PE^cKO^* CreERT2 iPSCs with (+) or without (-) TAM for 48h prior to collection and is used for differential gene expression and splicing analysis. **(B)** *PE* splicing and protein expression in *WT*, *SRSF3-PE^cKO^*, and *TRA2*β*-PE^cKO^*KOLF2.1J-CreERT2 iPSCs with (+) or without (-) TAM for 48h. *PE* splicing is measured by semi-quantitative RT-PCR using primers that amplify the included and skipped isoforms, and quantified as percent spliced-in (PSI). Cells are treated with mock or cycloheximide (+CHX) for 6h prior to collection to stabilize NMD-sensitive PE-containing transcripts. Protein expression is measured by Western blot, normalized to GAPDH loading control, and shown as fold change (FC) from +TAM to -TAM. Representative images are shown along with quantification (n=3, mean±SD; t-test +TAM to -TAM; \**P*<0.05, \*\**P*<0.01, \*\*\**P*<0.001, \*\*\*\**P*<0.0001). **(C,D)** Number of significant differential AS events (|ΔPSI_+TAM vs.-TAM_ |>10%, FDR<0.05) in *WT*, *SRSF3-PE^cKO^*, or *TRA2*β*-PE^cKO^*KOLF2.1J-CreERT2 (top) or WTC11-CreERT2 (bottom) iPSCs (n=3 per genotype and treatment) showing event type categories (A3SS: Alternative 3’ splice site; A5SS: Alternative 5’ splice site; CA: Cassette alternative exon; MXE: Mutually exclusive exons; RI: Retained introns) (**C**). ΔPSI distribution of significant AS events in *SRSF3-PE^cKO^* and *TRA2*β*-PE^cKO^*KOLF2.1J-CreERT2 (top) or WTC11-CreERT2 (bottom) iPSCs (**D**). TAM-induced AS events from +TAM WT iPSCs are removed from further analysis. **(E)** Significant differential AS events (|ΔPSI_+TAM vs.-TAM_ |>10%, FDR<0.05) overlapping across both cell lines (KOLF2.1J-CreERT2 and WTC11-CreERT2) and both genotypes (*SRSF3-PE^cKO^* and *TRA2*β*-PE^cKO^*) are grouped by more skipped across conditions, more included across all conditions, or divergent between genotypes. Gene enrichment analysis using GO biological processes was performed on spliced genes from each group. **(F-H)** A pgRNA CRISPR screen is used to delete shared skipped cassette exons from (**E**) and assay their impact on cell fitness in iCas9-WTC11 iPSCs **(F)**. Ranking of pgRNAs from the screen including positive controls (grey) and tested exons (blue), with significant hits highlighted (-Log_2_FC>1.2, *P_adj_*<0.05) **(G)**. Gene enrichment analysis using Reactome was performed using the top significant hits **(H)**. **(I)** Splicing in a cassette exon in *CLASP1* shown as schematic (top) is detected by RNA-seq data (middle panel) and validated by RT-PCR with primers that amplify the skipped and included isoforms (bottom panel) in KOLF2.1J-CreERT2 and WTC11-CreERT2 iPSCs (n=3 per genotype and treatment; mean±SD; \*\*\*\**P*<0.0001, \*\*\**P*<0.001, \*\**P*<0.01, ns – non-significant; two-tailed unpaired t-test). Computationally predicted functional consequences of included or skipped isoforms are indicated. **(J)** Splicing of the *CLASP1* cassette exon is measured by RT-PCR with primers that amplify the skipped and included isoforms in iCas9-WTC11 iPSCs expressing pgRNAs targeting the *CLASP1* cassette exon with (+) and without (-) DOX (n=3; mean±SD; \*\*\*\**P*<0.0001 two-tailed unpaired t-test). (**K,L**) Relative cell numbers in iCas9-WTC11 iPSCs expressing pgRNAs targeting the *CLASP1* cassette exon, or HNRNPL as a positive control, or AAVS as negative control at indicated timepoints following Cas9 activation (n=3; mean±SD; \*\*\**P* 0.001, two-way ANNOVA with Tukey correction to pgAAVS) **(K)**. Representative phase contrast images showing the morphology of iPSCs cells from (k) at indicated timepoints following Cas9 activation **(L)**.

### PE deletions compromise iPSC fitness and early progenitor viability without biasing lineage specification

To determine whether PE essentiality extends beyond the pluripotent state to actively cycling progenitors, and whether PE deletion influences early lineage specification, we next examined how PE deletion affects early differentiation capacity. To test whether PE deletion biases lineage commitment, we cultured *WT*, *SRSF3-PE^cKO^* and *TRA2*β*-PE^cKO^* iPSC in non-conditioned media permitting spontaneous differentiation, with staining for iPSC and early lineage markers at days 5 and 7 post-TAM (**Fig.S3A**). Despite baseline variability in spontaneous differentiation efficiency among clones, TAM treatment did not induce differentiation or skew cells toward any germ layer but did produce a marked reduction in cell viability for *SRSF3-PE^cKO^* and *TRA2*β*-PE^cKO^*, but not *WT* lines across all conditions (**Fig.S3A**). This confirms that PE-deletion lethality is not an artifact of stem cell maintenance media and extends to actively cycling progenitor and transitional cell states. Consistent with this, PE-deleted and OCTA4A^+^ iPSCs displayed pronounced abnormalities in nuclear morphology, including a ∼2-fold increase in nuclear area with significant asymmetry, marked by reduced nuclear roundness and width-to-length ratio (**Fig. S3B-C**), phenotypes potentially linked to defects in genome segregation during mitosis.

Similarly, when PE deletions were induced on day 1 of directed differentiation toward neuroectodermal, mesodermal, or endodermal progenitors using lineage-specific preconditioned media, TAM-treated *PE^cKO^*, but no *WT*, cells showed a global decrease in total cell number across all three lineage conditions (**Fig.S3D**), consistent with the proliferative lethality described above. In this setting, we did observe differences in early lineage differentiation, primarily a reduction in SOX17+ endoderm progenitors in both *SRSF3-PE^cKO^* and *TRA2*β*-PE^cKO^*lines, and reduction in SOX1+ neuroectoderm cells in *SRSF3-PE^cKO^*lines. Brachyury+ mesoderm progenitors were not different between treatment conditions (**Fig.S3D**), overall, suggesting that PE deletion does not completely eliminate lineage differentiation potential, but may influence differentiation efficiency and the proliferative capacity and viability of iPSCs and early progenitor cells.

### PE deletions trigger convergent, widespread splicing dysregulation and disrupt splicing-factor cross-regulatory networks in iPSCs

Having established that PE deletion is lethal across pluripotent and early progenitor states, we next sought to define the molecular basis of this requirement by characterizing the transcriptomic and splicing consequences of PE loss in iPSCs. We performed high-depth RNA-sequencing (RNA-seq) in *WT*, *SRSF3-PE^cKO^,* and *TRA2*β*-PE^cKO^*KOLF2.1J and WTC11 iPSC lines at 48 hours post-TAM, a timepoint at which PE deletion is complete and protein upregulation is established, but before apoptotic cell death dominates the culture (**Fig.2A-B**; **Fig.S2**). TAM treatment of WT iPSCs resulted in fewer than 50 differentially expressed genes (DEGs, |Log_2_FC|>1, *P_adj_*<0.05) and minimal splicing changes, confirming that TAM itself does not substantially perturb the transcriptome (**Fig.2C-D**; **Fig.S4A**; **Tables S1-2**).

PE deletion produced extensive transcriptomic reorganization in both genotypes, with effects detectable across both broadly expressed and pluripotency-associated gene sets. *SRSF3-PE^cKO^* iPSCs exhibited 246 and 651 upregulated and 406 and 825 downregulated genes in KOLF2.1J and WTC11, respectively; while *TRA2*β*-PE^cKO^* iPSCs showed 650 and 710 upregulated and 1,499 and 2,229 downregulated genes (**Fig.S4A-B**; **Table S1**). To assess whether PE deletion affects the pluripotent transcriptional state, we examined a curated 12 marker genes signature derived from well-established stem cell consortia studies that monitor the activation of the core transcriptional network and the suppression of differentiation-associated pathways (*47, 48*). While several of these genes exhibited significant differential expression, none were overtly up- or down-regulated (|Log_2_FC|>1). Additionally, only three exhibited AS changes upon TAM treatment in at least one cell line and genotype (**Fig.S4C**). This suggests a modest perturbation of the pluripotent transcriptional state rather than a directed differentiation response. We next investigated the splicing response using rMATS to quantify event-level splicing with a “Percent Spliced-In” (PSI) value for each AS event (*8–10, 49*), identifying 6,771 and 12,870 TAM-induced significant AS events in *SRSF3-PE^cKO^*and *TRA2*β*-PE^cKO^*KOLF2.1J iPSCs, and 8,785 and 12,687 events in the corresponding WTC11 lines (**Fig.2C-D**; **Table S2**). Splicing profiles of TAM-treated PE^cKO^ iPSCs were clearly separated from untreated controls and from each other by principal component analysis, indicating that SRSF3 and TRA2β regulate largely distinct, though partially overlapping, splicing programs (**Fig.S5A**). Across both cell lines and genotypes, ∼75% of significant AS events were computationally predicted to alter known amino acid composition, disrupt known protein domains, or introduce premature termination codons that target the transcript for NMD, suggesting broad functional impact of PE deletions on the iPSC proteome (**Fig.S5B**).

The most informative finding within this dataset is the degree of cross-genotype and cross-line convergence. Of all AS events, 3,725 were shared in *SRSF3-PE^cKO^* KOLF2.1J and WTC11, and 6,808 between *TRA2*β*-PE^cKO^* KOLF2.1J and WTC11 — all regulated in the same direction, identifying a cell-line-agnostic splicing response for each factor (**Fig.S5C**; **Table S2**). Critically, 2,326 and 2,965 AS events were shared between *SRSF3-PE^cKO^* and *TRA2*β*-PE^cKO^* within KOLF2.1J and WTC11 lines, respectively (*P_adj_*<0.001) (**Fig.S5D**; **Table S2**). Of these shared events, approximately two-thirds were regulated concordantly; the remaining third were regulated in opposing directions, indicating that SRSF3 and TRA2β act as both cooperative and antagonistic regulators at a subset of shared targets.

Intersecting all four conditions (both genotypes, both lines) identified 1,114 high-confidence shared AS events (**Fig. 2E**; **Table S3**). Among these, 342 were shared and skipped, enriched in genes governing embryonic and organismal development; 355 were shared and included, enriched in microtubule-based processes and DNA repair; and 417 showed opposing directionality between genotypes, enriched in membraneless organelle assembly and RNA processing (**Fig.2E**; **Table S3**). Gene ontology enrichment of spliced genes across all four conditions independently converged on DNA repair, mitosis, and microtubule organization as top pathways (**Fig.S5E**; **Table S2**). We validated representative AS events by RT-PCR in both iPSC lines, including splicing changes in histone regulators (*ARID1B, BRD8, EZH2, SIRT1, PRDM5*), RNA splicing factors (*RBM39, ESRP1*), DNA damage response factors (*ATR*) and pluripotency regulators *(TCF3, ESRP1*) (**Fig.S6**). Notably, both genotypes exhibited decreased intron retention in the 3’UTR of *TCF3*, a transcriptional repressor of self-renewal (*50–52*), and opposing cassette exon changes in *ESRP1*, a critical regulator of *OCT4*, *NANOG*, and *SOX2* (*53, 54*) (**Fig.S6**), suggesting that while core pluripotency factors were not significantly changing (**Fig.S4C**), PE-deletion was impacting accessory regulators of pluripotency.

To distinguish direct splicing effects from those propagated through altered levels of other SFs, we mapped experimentally-derived SRSF3 and TRA2β binding motifs (*55–57*) across all significant cassette exon AS events, using a set of non-significant events as background to predict motif enrichment. SRSF3 motifs were enriched within exons showing increased inclusion in *SRSF3-PE^cKO^* iPSCs, and TRA2β motifs within those showing increased inclusion in *TRA2*β*-PE^cKO^* iPSCs (**Fig.S7A-B**), consistent with the established role of both factors as activators of exon inclusion (*5, 58*), and confirming a direct component to the splicing response.

Beyond direct effects, PE deletion disrupted the broader splicing factor regulatory network. Among 369 RNA-binding proteins (RBPs) surveyed, including SR proteins, hnRNPs, and spliceosome components, 104 were differentially expressed and 229 AS events were differentially spliced across datasets, with only 16 DEGs and 42 AS events shared across both genotypes and both lines (**Fig.S7C-E**; **Table S4**). The limited overlap between differentially expressed and differentially spliced RBPs indicates that transcriptional and post-transcriptional cross-regulation operate in parallel rather than in series. Among validated cross-regulatory events, *TRA2*β*-PE^cKO^* induced 100% inclusion of the *TRA2*α*-PE*, correlating with ∼90% reduction in TRA2α protein (**Fig.S7F**), a cross-regulatory relationship previously described in somatic cell types (*8, 9, 58*). Overall, PE deletions in *SRSF3* and *TRA2*β each triggered distinct downstream RBP changes indicating that while both PEs feed into shared splicing programs, they do so through non-identical regulatory hierarchies.

### A CRISPR exon-deletion screen identifies mitotic regulators as functional drivers of PE-deletion lethality

The convergence of 1,114 shared AS events across both genotypes and iPSC lines raised the question of which specific splicing changes drive cell death, as opposed to reflecting downstream transcriptomic noise. Here, we focused on skipped exons because their deletion can be modeled efficiently using paired guide RNAs (pgRNAs) targeting individual exons (*13*), enabling high-throughput functional interrogation, an approach that is not directly applicable to included exons or more complex AS event types. We designed a pooled pgRNAs CRISPR screen targeting 274 shared cassette exons that were skipped in both *SRSF3-PE^cKO^*and *TRA2*β*-PE^cKO^*(**Fig.2F**). Each exon was targeted with 9 pgRNAs, yielding a library of ∼2,186 pgRNAs, including 75 non-targeting controls and 153 positive controls targeting known essential exons for 17 different proteins. The screen was conducted in doxycycline-inducible Cas9 (iCas9) WTC11 iPSCs across four independent replicates, with pgRNA representation quantified by deep sequencing at day 0 and day 20 (**Fig.2F; Fig.S8A-C**). pgRNA representation at day 0 correlated strongly with the input plasmid pool (r_2_>0.9), and replicate concordance was high at all timepoints (**Fig.S8C**).

As expected, pgRNAs targeting essential genes were significantly depleted by day 20, while the majority of candidate exon deletions had no measurable effect on iPSC fitness (**Fig.2G; Fig.S8D-E**). Applying cutoffs designed to ensure high sensitivity while capturing all positive control targets (log_2_FC<-1.2 and *P*_adj_<0.05), the screen identified 19 cassette exon deletions that significantly reduced cell fitness (**Fig.2G**; **Table S5**). Top-ranking fitness hits were enriched in genes regulating cell cycle progression and the G2/M transition (**Fig.2H**; **Table S5**), consistent with a role for SRSF3- and TRA2β-regulated splicing of mitotic genes as a functional contributor to iPSC lethality.

The highest-confidence individual hit in this category was a cassette exon in *CLASP1* (cytoplasmic linker-associated protein 1), a core component of the mitotic spindle apparatus required for kinetochore–microtubule attachment and chromosome alignment (*59–61*). RT-PCR confirmed that *CLASP1* exon inclusion decreases in TAM-treated *SRSF3-PE^cKO^* and *TRA2*β*-PE^cKO^* iPSCs (**Fig.2I**); as this exon is present in a non-canonical isoform, its skipping is predicted to shift expression toward the canonical full-length CLASP1 protein. Validations using individual two pgRNAs confirmed efficient exon deletion and a reproducible reduction in iPSC fitness (**Fig.2J-L; Fig.S8F**), supporting *CLASP1* exon skipping as a functional contributor to the PE-deletion phenotype. Consistent with a functional consequence of this isoform switch, AlphaFold structural modeling predicts differences in the three-dimensional conformation of CLASP1 protein between the exon-included and exon-skipped isoforms (**Fig.S8G**). Additional screen hits included cassette exons in *SPATS2*, an RBP linked to cell-cycle regulation and cancer progression (*62*); *PHYKPL*, a nuclear-encoded mitochondrial metabolic gene, and *APBB2*, an adaptor protein at the intersection of APP signaling, neurodegeneration, and cell cycle control (*63*), all of which showed concordant splicing changes in both *PE^cKO^* genotypes (**Fig.S8H**).

### PE deletions induce mitotic spindle defects across a shared program of cell-cycle-associated splicing changes

Beyond *CLASP1*, our transcriptomic analysis identified 31 additional significant AS events across 21 mitosis-associated genes shared between TAM-treated *SRSF3-PE^cKO^* and *TRA2*β*-PE^cKO^* iPSCs in both KOLF2.1J and WTC11 lines (**Fig.3A; Table S6**). These genes span three functional modules of mitotic fidelity: i) pre-mitotic genome integrity and checkpoint control (*DBF4B*, *TAF2*, *RNASEH2B*, *BABAM2*, *PLK3*, *CDC14B*); ii) centrosome duplication, spindle assembly, and chromosome alignment (*STIL*, *OFD1*, *NDE1*, *TACC1*, *TACC2*, *EML1*, *CLASP1*, *ZNF207*, *CENPE*, *BIRC5*); and iii) cytokinesis and post-mitotic architecture (*KIF23*, *BIRC5*, *DNM2*, *GOLGA2*, *NUP62*). The convergence of splicing dysregulation across all three modules suggests a broad impairment of mitotic fidelity rather than disruption of a single pathway.

**Figure 3:**
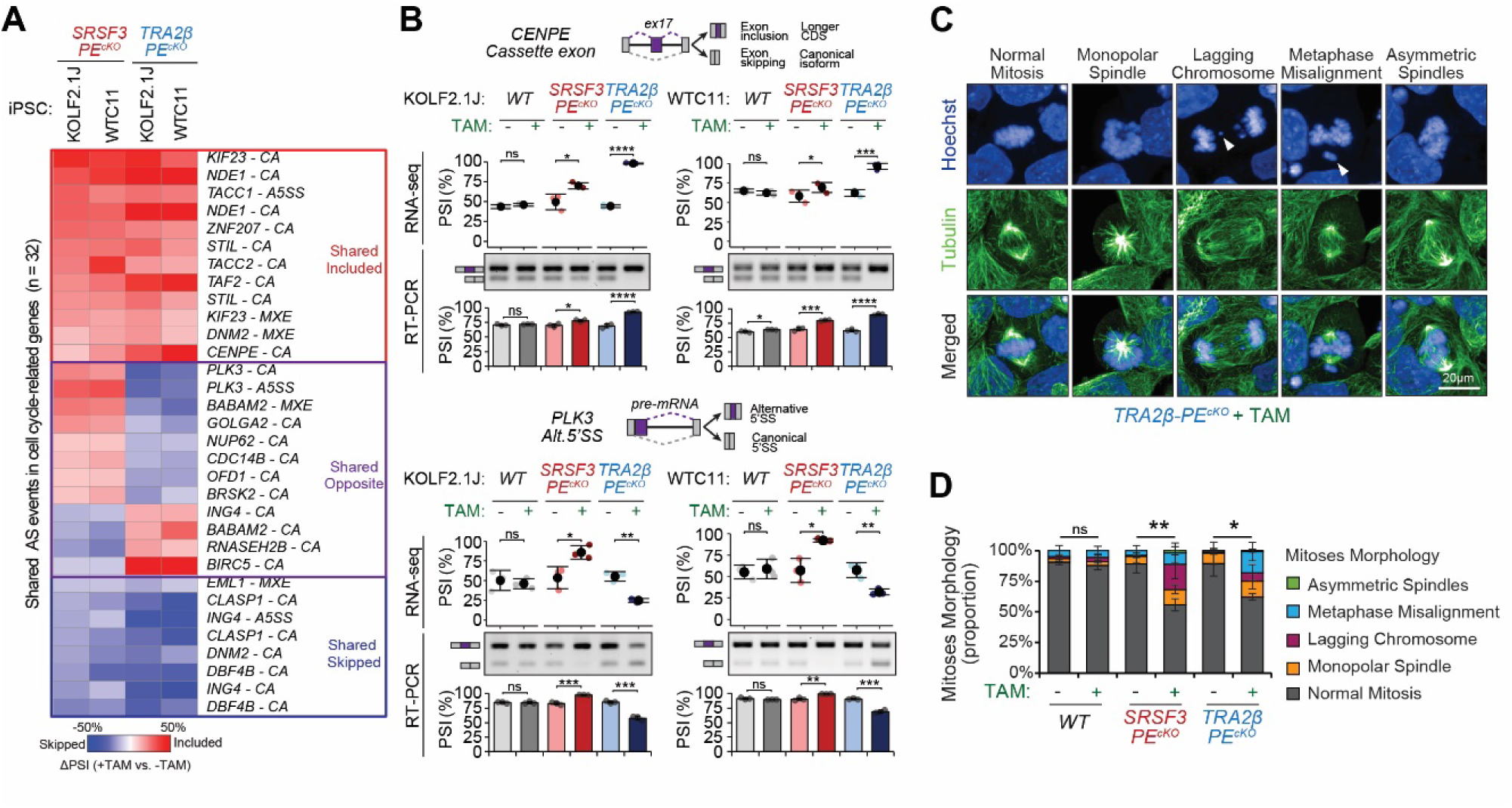
*SRSF3-PE* and *TRA2*β*-PE* deletions causes abnormal splicing of mitosis-related transcripts and induces abnormalities in iPSC mitosis. **(A)** Significant differential AS events (|ΔPSI_+TAM vs.-TAM_ |>10%, FDR<0.05) in mitosis-related genes overlapping across both cell lines (KOLF2.1J-CreERT2 and WTC11-CreERT2) and both genotypes (*SRSF3-PE^cKO^* and *TRA2*β*-PE^cKO^*) are grouped by more skipped across conditions, more included across all conditions, or divergent between genotypes. **(B)** Examples of spliced mitosis related genes shown as schematic (top) is detected by RNA-seq data (middle panel) and validated by RT-PCR with primers that amplify the skipped and included isoforms (bottom panel) in KOLF2.1J-CreERT2 and WTC11-CreERT2 iPSCs (n=3 per genotype and treatment; mean±SD; \*\*\*\**P*<0.0001, \*\*\**P*<0.001, \*\**P*<0.01, ns – non-significant; two-tailed unpaired t-test). AS event types and computationally predicted functional consequences of included or skipped isoforms are indicated. **(C,D)** Representative images of mitosis defects in TAM-treated *TRA2*β*-PE^cKO^*KOLF2.1J iPSCs detected using immunofluorescence with tubulin as a microtubule marker and Hoechst as a DNA marker (**C**). Normal mitoses and mitotic defects were quantified in *WT*, *SRSF3-PE^cKO^*, and *TRA2*β*-PE^cKO^* KOLF2.1J iPSCs at 48h following TAM treatment (**D**). Cells were treated with nocodazole for 12h, followed by withdrawal for 2h prior to fixation, in order to enrich for cells actively undergoing mitosis. Mitoses morphology is represented as proportion of normal mitoses, monopolar spindles, lagging chromosomes, metaphase misalignment, and asymmetric spindles compared to total mitoses (n=3 per genotype and treatment with average of 36 mitotic cells analyzed per replicate; mean±SD; \*\*\*\**P*<0.0001, \*\*\**P*<0.001, \*\**P*<0.01, \**P*<0.05, ns – non-significant; two-tailed unpaired t-test).

Of the 32 total shared AS events, 20 were regulated in the same direction across all four conditions, including *CLASP1* and four additional cassette exons that were either screened but did not significantly decrease cell fitness, or were not amenable to pgRNA targeting. The remaining 12 shared events were regulated in opposing directions between *SRSF3-PE^cKO^*and *TRA2*β*-PE^cKO^*. For example, a *CENPE* cassette exon showed increased inclusion upon both PE deletions (**Fig.3B**). Conversely, *SRSF3-PE* deletion promoted increased alternative 5’SS usage in *PLK3*, whereas *TRA2*β*-PE* deletion produced the opposite pattern (**Fig.3B**). These bidirectional effects are consistent with the known antagonistic activities of SRSF3 and TRA2β at shared targets and suggest that the net functional outcome at individual mitotic genes depends on the relative stoichiometry of both factors.

To determine whether these splicing alterations produce measurable mitotic defects, we quantified mitotic abnormalities in WT, *SRSF3-PE^cKO^*, and *TRA2*β*-PE^cKO^* KOLF2.1J iPSCs at 48 hours post-TAM, following a 12-hour nocodazole synchronization and release to enrich for actively dividing cells. (**Fig.3C-D**). Mitotic states were classified by immunofluorescence as normal or defective based on microtubule spindle and chromosome morphology and expressed as a proportion of total mitoses (**Fig.3C**). Both *SRSF3*-*PE^cKO^* and *TRA2*β-*PE^cKO^* cells showed a significant increase in abnormal mitoses relative to TAM-treated WT and untreated controls, with elevated frequencies of monopolar spindles, lagging chromosomes, and metaphase misalignment (**Fig.3D**). These mitotic phenotypes directly parallel the functional categories of splicing-dysregulated genes identified above, supporting a model in which convergent PE-dependent splicing programs are required to maintain the fidelity of cell division in proliferating stem cells.

### *TRA2β-PE* deletion is tolerated in post-mitotic neurons, revealing a proliferation-dependent requirement

The mitotic specificity of these defects raised a fundamental question: whether PE essentiality is a general property of all cell types or restricted to actively dividing cells. Given the well-established role of *TRA2*β in brain development (*32, 33*), we asked whether post-mitotic neurons could tolerate *TRA2*β*-PE* deletion. We induced *TRA2*β*-PE* deletion in fully differentiated post-mitotic neurons generated from KOLF2.1J iPSCs via doxycycline-inducible *NEUROG2* (iNGN2) overexpression, a protocol that produces mature excitatory neurons within 30 days (*64*). TAM was administered at day 10 of differentiation to avoid early progenitor death (**Fig.4A**). *TRA2*β*-PE* deletion was confirmed at both RNA and protein levels, with complete loss of PE inclusion and a ∼2.5-fold increase in TRA2β protein (**Fig.4B**). In contrast to iPSCs, PE deletion was well tolerated: TAM-treated TRA2β-PE^cKO^ neurons maintained normal morphology and expressed neuronal markers (NeuN and MAP2) at levels comparable to untreated and WT controls (**Fig.4C-D**; **Fig.S9A**). No KI67^+^ proliferative progenitors were detected under standard mitotic inhibitor conditions in either genotype (**Fig.S9A**), consistent with a fully post-mitotic culture.

**Figure 4:**
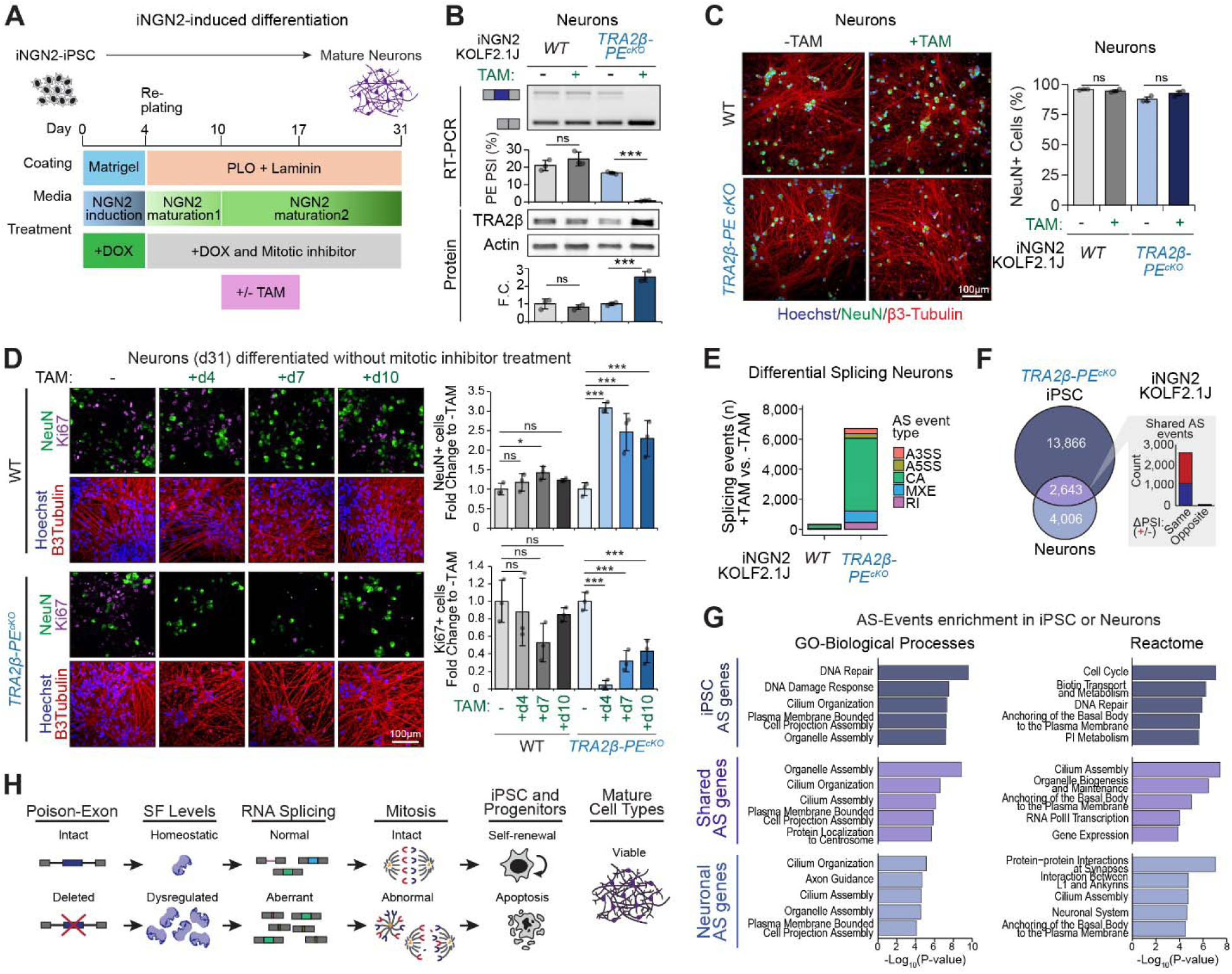
*TRA2*β*-PE* deletion is tolerated in mature post-mitotic cell types. **(A)** Schematic of the KOLF2.1J-iNGN2 iPSCs neuronal differentiation protocol. DOX treatment induces NGN2 expression, while TAM treatment induces PE deletion. **(B)** *PE* splicing and protein expression in *WT*, and *TRA2*β*-PE^cKO^*KOLF2.1J iNGN2 CreERT2 iPSC-derived neurons at d31 with (+) or without (-) TAM. *PE* splicing is measured by semi-quantitative RT-PCR using primers that amplify the included and skipped isoforms, and quantified as percent spliced-in (PSI). Protein expression is measured by Western blot, normalized to Actin loading control, and shown as fold change (FC) from +TAM to -TAM. Representative images are shown along with quantification (n=3, mean±SD; t-test +TAM to -TAM; \**P*<0.05, \*\**P*<0.01, \*\*\**P*<0.001, \*\*\*\**P*<0.0001). **(C)** *WT* and *TRA2*β*-PE^cKO^*iNGN2 CreERT2 iPSC-derived neurons were stained at day 31 for neuronal markers (NeuN and β3-tubulin) and counterstained with Hoechst. Neuronal cell proportion is estimated as the percentage of NeuN+ cells (n=3, mean±SD; t-test +TAM to -TAM; \**P*<0.05, ns: not-significant). **(D)** *WT* and *TRA2*β*-PE^cKO^* iPSC-derived neurons differentiated without a mitotic inhibitor were stained at day 31 for the neuronal markers NeuN and β3-tubulin, progenitor proliferative marker Ki67, and counterstained with Hoechst. Neuronal cell proportion (NeuN+) and proliferative cell proportions (Ki67+) are quantified and shown as fold change to -TAM (n=3, mean±SD; t-test +TAM to -TAM; \**P*<0.05, ns: not-significant). **(E)** Number of significant differential AS events (|ΔPSI_+TAM vs.-TAM_ |>10%, FDR<0.05) in *WT* or *TRA2*β*-PE^cKO^*KOLF2.1J-iNGN2 CreERT2 iPSC-derived neurons at d31 (n=3 per genotype and treatment) showing event type categories (A3SS: Alternative 3’ splice site; A5SS: Alternative 5’ splice site; CA: Cassette alternative exon; MXE: Mutually exclusive exons; RI: Retained introns) TAM-induced AS events from +TAM WT neurons are removed from further analysis. **(F,G)** Number of unique and shared significant AS events (|ΔPSI_+TAM vs.-TAM_ |>10%, FDR<0.05) between *TRA2*β*-PE^cKO^*KOLF2.1J iNGN2 CreERT2 iPSCs and iPSC-derived neurons (*P_adj_*<0.001; Chi-square with Monte-Carlo adjustment) **(F)**. ΔPSI directionality of shared AS events is plotted on the right (same: ΔPSI<0 (blue) or ΔPSI>0 (red) in both cell lines; opposite: ΔPSI<0 in one cell line while ΔPSI>0 in the other). Gene enrichment analysis using GO biological processes and Reactome terms was performed on spliced genes unique to *TRA2*β*-PE^cKO^* iPSC, shared between iPSC and neurons, or unique to neurons, with shown top significantly enriched pathways at *p_adj_*<0.05 **(G)**. **(H)** Graphical abstract: SR proteins SRSF3 and TRA2β control their own expression through ultraconserved PEs that trigger NMD and their deletion elevates SR protein levels. In proliferating iPSCs, PE deletion disrupts the splicing of mitotic regulators, causing spindle defects and apoptotic cell death, whereas in post-mitotic neurons PE deletion is well tolerated despite widespread splicing changes in genes regulating synaptic function.

To directly test whether the survival advantage in neurons reflects their post-mitotic state, we repeated the experiment after withdrawing mitotic inhibitors, allowing residual proliferative cells to persist. Under these conditions, TAM-induced *TRA2*β*-PE* deletion produced a drastic reduction in KI67^+^ cells and a corresponding 2- to 3-fold increase in proportion of NeuN^+^ neurons, with the strongest effect when TAM was administered on day 4 (**Fig.4D**). Increased cellular debris was evident by β-tubulin and Hoechst staining, consistent with selective death of cycling cells (**Fig.4D**). This establishes that *TRA2*β*-PE* is dispensable in NGN2-derived post-mitotic neurons but essential in proliferating cells within the same differentiation context.

Despite preserved viability, RNA-seq revealed that *TRA2*β-*PE^cKO^* neurons displayed a substantial transcriptomic response. TAM treatment of *WT* neurons produced minimal changes, while *TRA2*β*-PE* deleted neurons exhibited 931 upregulated and 793 downregulated genes (**Fig.S9C**; **Table S7**). Critically, both WT and PE-deleted neurons showed equivalent downregulation of iPSC markers and upregulation of mature neuronal markers (**Fig.S9D**), confirming successful and equivalent differentiation. Direct comparison of *WT* and *TRA2*β*-PE^cKO^* neurons revealed largely modest changes in known iPSC, pan-neuronal, and subtype-specific markers, with only *LEFTY1* (an iPSC marker) and *PNMT* (a marker of adrenergic neurons) significantly altered (|log_2_FC|>1, *P*_adj_<0.05) (**Fig.S9E**), overall reflective of modest perturbation of neuronal identity that is distinct from the catastrophic transcriptomic disruption seen in iPSCs.

At the splicing level, *PE-*deleted neurons exhibited 6,649 AS events (**Fig.4E**; **Table S8**) of which ∼80% are computationally predicted to alter known amino acid composition, disrupt known protein domains, or introduce premature termination codons that target the transcript for NMD — suggesting broad functional impact on the neuronal proteome (**Fig.S9F**). Of these 2,643 were shared with iPSCs and largely regulated in the same direction (**Fig.4F**). TRA2β binding motifs were enriched in differentially spliced cassette exons in neurons as in iPSCs (**Fig.S10A**), confirming direct regulatory continuity across cell types. As in iPSCs, *TRA2*β*-PE* deletion altered expression and splicing of other RBPs in neurons, though the breadth of cross-regulatory changes was markedly reduced compared to iPSCs (**Fig.S10B-D**; **Table S9**). In contrast to iPSCs, the vast majority of AS events in mitosis-related genes detected upon PE deletion in iPSCs were not significantly altered in neurons, consistent with the post-mitotic state of the iNGN2 culture (**Fig.S9H**). Instead, the AS events detected only in *TRA2*β*-PE* deleted neurons were enriched in genes involved in cilium assembly, cell projection, and synaptic plasticity (**Fig.4G**). Validated neuron-specific events converged on genes critical to excitatory synaptic function including: i) axonal output with an intron retention event in *ANK3*, which encodes Ankyrin-G, the master scaffold of the axon initial segment required for sodium channel clustering and action potential initiation (*65*); ii) presynaptic vesicle release with increased skipping of a cassette exon in *RIMS2*, which encodes a core active zone protein that tethers calcium channels to release sites and primes synaptic vesicles for exocytosis (*66*); iii) vesicle trafficking and recycling with increased inclusion of a cassette exon in *DENND5A*, a Rab-GEF predominantly expressed in neuronal tissues whose loss-of-function mutations cause epileptic encephalopathy (*67*); and iv) synaptic RNA regulation with increased cassette exon inclusion in *FXR1,* implicated in synapse homeostasis (*68, 69*) (**Fig.S9G**). The dissociation between extensive splicing changes and preserved viability in neurons underscores that splicing dysregulation is not sufficient for lethality, rather its link to a mitotic context is required. The neuron-specific splicing program controlled by *TRA2*β*-PE* instead points to a post-mitotic function in synaptic plasticity and neuronal architecture that warrants dedicated investigation.

## Discussion

Poison exon-mediated AS-NMD has long been recognized as a homeostatic mechanism for controlling splicing factor expression, yet its biological necessity — when it matters, in which cell types, and through which downstream effectors — has remained poorly defined. The data presented here addresses these questions for two PEs embedded in ultraconserved genomic elements in SR protein genes. Homozygous deletion of *SRSF3*-*PE* or *TRA2*β-*PE* is selected against in mouse embryos and human iPSCs, indicating that PE-mediated dosage control is under strong selection pressure from the earliest stages of mammalian development. Conditional PE deletion in iPSCs causes apoptotic cell death while remaining tolerated in post-mitotic neurons, mapping PE essentiality to proliferative state rather than lineage identity. Together, these observations support a model in which precise SR protein dosage, enforced through PE-mediated AS-NMD, is required for the fidelity of mitotic splicing programs in proliferating stem cells.

The AS-NMD mechanism that couples PE inclusion to SR protein degradation has been established as a general autoregulatory feedback loop (*7–9, 15, 16*), but why this mechanism requires such extreme sequence conservation, and in which contexts its disruption is consequential, has been unknown. Our data suggests that PE-mediated dosage control is dispensable in post-mitotic cells, where transcriptomic demands on splicing fidelity are relatively static, but is acutely essential in proliferating cells, where a large and dynamic set of mitosis-associated AS events must be precisely coordinated across each cell cycle. Both too little SR protein, as occurs when PEs are aberrantly over-included, and too much, as occurs when PE deletion abrogates AS-NMD, are incompatible with normal division. This suggests a possible “goldilocks” model paralleled with cancer biology, where *SRSF3* and *TRA2*β are elevated as oncoproteins and forced PE inclusion impairs tumor cell viability (*8, 9, 13*), and with our finding that PE deletion is equally lethal to actively cycling pluripotent and lineage progenitor cells.

Further evidence to this hypothesis comes from the recent demonstration that deletion of the ultraconserved *Tra2b*-PE in mice causes azoospermia through catastrophic cell death during meiotic prophase, driven by aberrant TRA2β accumulation (*70*). The parallel lethality of *Tra2b*-PE deletion in both meiotic and mitotic division, across independent biological systems, reinforces the conclusion that precise PE-mediated dosage control is a conserved and essential component of the cell division machinery. The discrepancy between that study’s finding that early mitotically active spermatogonia are spared, and our finding of mitotic sensitivity in iPSCs, most likely reflects cell-type-specific differences in downstream TRA2β splicing targets rather than a fundamental mechanistic difference.

The identification of mitotic regulators as the primary functional targets of PE-controlled splicing connects to an emerging literature on the coordination between the splicing apparatus and cell division. SR proteins are known to dissociate from hyperphosphorylated mitotic chromosomes in an Aurora B kinase-dependent manner and reassociate during mitotic exit (*71*), further Aurora A directly phosphorylates SR proteins, including SRSF3, indirectly regulating AS of hundreds of cell-cycle genes (*72*). Our data extend this model: PE-dependent control of SRSF3 and TRA2β levels is required to maintain correct AS of a broad program of mitosis-associated genes, and its disruption is associated with spindle defects, lagging chromosomes, and metaphase misalignment. The *CLASP1* AS event, a high-confidence functional hit in our CRISPR screen, provides one concrete molecular link between PE-controlled splicing and mitotic fidelity, though the overall lethality phenotype most likely reflects the cumulative disruption of multiple AS events rather than the consequence of any single isoform switch.

A notable complexity in our data is that, while *SRSF3-PE* and *TRA2*β*-PE* deletions produce convergent cellular phenotypes, their downstream splicing programs show both extensive overlap and systematic antagonism, and resolving how SRSF3 and TRA2β regulate overlapping but distinct and partially antagonistic mitotic splicing programs will be critical to understanding the full mechanistic picture. Approximately one-third of shared AS events are regulated in opposing directions at loci including *CENPE*, *PLK3*, and *AURKB*. This bidirectionality is consistent with known biochemical antagonism between splicing factors at overlapping binding sites (*9, 10, 73, 74*), and suggests that the net functional outcome at individual mitotic genes depends on the relative stoichiometry of both factors simultaneously.

The resolution of the UCE conservation paradox that emerges from this work is mechanistically distinct from prior explanations based on enhancer function or RNA structural constraints. UCE-embedded PEs are conserved because any sequence variation that alters PE splicing efficiency would perturb SR protein dosage sufficiently to compromise mitotic fidelity in proliferating cells. Unlike enhancers, which can tolerate compensatory mutations within their binding site sequences (*4*), the *SRSF3-PE* and *TRA2*β*-PE* regulatory elements do not appear to accommodate sequence drift without detriment to cell fitness. Consistent with this, systematic deletion mutagenesis and dCasRx scanning across the *TRA2*β-*PE* UCE demonstrated that every position within the element is required for normal PE splicing regulation, with no dispensable sequence windows detectable (*9*). The cryptic 3’SS rescue in homozygous *SRSF3-PE-3’SS^mut/mut^*iPSC clones further reinforces this constraint: when the canonical mechanism is destroyed, single human iPSC clones that survive homozygous disruption do so by regenerating PE function *de novo*, underscoring the irreplaceable nature of this regulatory axis.

Beyond stem cell biology, PE-mediated AS-NMD of SR proteins is dysregulated across a broad spectrum of human diseases. In cancer, *SRSF3-PE* and *TRA2*β-*PE* are preferentially skipped relative to normal tissue, enabling oncogenic SR protein overexpression; splice-switching ASOs restoring *TRA2*β*-PE* inclusion impair cancer cell viability and induce expression of a PE-containing lncRNA that sequesters nuclear proteins and amplifies anti-tumor effects in organoid and patient-derived xenograft models (*8, 9*). Additionally, recent work has described recurrent mutations in a subset of UCEs, including micro-RNA loci that are regulated by SRSF3, underscoring complex interactions between different types of UCEs in disease states (*75*). Beyond cancer, *TRA2*β*-PE* skipping gates TCR sensitivity and effector T cell expansion during antigen stimulation, while PE re-inclusion enables T cell survival as antigen wanes (*7*), and SR protein splicing dysregulation has been implicated in cardiac injury and neurodegeneration (*5, 24, 27, 35, 76*). PE-targeting strategies are mechanistically attractive precisely because tumor cells that have already downregulated PE inclusion are more sensitive to forced PE re-inclusion than normal cells in which PE is physiologically maintained (*8, 9*), an intrinsic tumor selectivity that the present study refines further. Because PE essentiality is mitotic-state-dependent, on-target toxicity in normal tissues is predicted to be concentrated in rapidly dividing progenitor compartments, while post-mitotic tissues are largely spared — providing a clear rationale for tumor-targeted delivery modalities such as tumor-selective ASO formulations or antibody-conjugated splice-switching oligonucleotides. The proliferation-dependence of PE essentiality is therefore not a barrier but an exploitable biological feature that aligns the most sensitive normal cell population with tissues amenable to sparing through targeted delivery.

A key question raised by these findings is whether other UCE-embedded PEs in splicing factors share this proliferative essentiality or whether the severity of PE loss is determined by the identity of the downstream splicing targets controlled by each factor. *SRSF3* and *TRA2*β are not unique in harboring deeply conserved PEs as each SR protein gene encodes at least one ultraconserved or deeply conserved PE region, as well as other splicing factors including members of the hnRNP family (*9, 15, 21, 77–81*). However, not all conserved PE deletions produce equivalent phenotypes. Belleville et al. recently showed that deletion of a conserved PE in *Smndc1*, a non-SR splicing factor, causes pervasive mRNA processing alterations and a transient pre-weaning growth restriction that normalizes after weaning in mice, but not lethality (*3*). This contrast with the acute iPSC lethality we observe for *SRSF3-PE* and *TRA2*β*-PE* suggests that PE conservation reflects the importance of precise dosage control for each individual factor. Systematic conditional deletion of PEs across the SR protein family, using the framework established here, would directly test whether mitotic splicing vulnerability is a general property of SR protein PEs or a specific feature of SRSF3 and TRA2β biology. More broadly, several questions raised by the present work define the agenda for future investigation: the mechanistic basis for the asymmetric lineage sensitivity of progenitors to PE deletion; the relative quantitative contribution of individual mitotic AS events to iPSC lethality; and whether the diffuse transcriptomic changes in *TRA2*β-*PE*-deleted neurons, including splicing alterations at synaptic loci, phenotypically manifest under conditions of neuronal stress or aging. These questions position UCE-embedded PE splicing as a regulatory axis with broad implications for stem cell biology, development, and disease.

## Acknowledgments

We thank all members of the Anczuków, Murray and Skarnes labs for helpful discussions and comments on the manuscript. We acknowledge assistance from Microscopy, Genome Editing, and Genome Technologies Sequencing Services at The Jackson Laboratory (JAX) supported in part by NCI grant P30CA034196. We also acknowledge the use of ChatGPT and Claude Code as assistive editing tools used to optimize code design and condense language during the editing process.

## Author contributions

Conceptualization: NL, MB, OA; Methodology: NL, MB, OA, IW, RE, JMD, WS, SM; Investigation: NL, MB, OA, IW, RE, MR, CH, JMD; Visualization: NL, MB, OA, IW, RE; Funding acquisition: OA; Project administration: OA; Supervision: OA; Writing – original draft: NL, MB, OA; Writing – review & editing: NL, MB, OA, IW, RE, MR, CH, JMD, WS, SM.

## Data, code and materials availability

Our splicing analysis pipeline v2.0 (*8–10*) is available on Github (https://github.com/TheJacksonLaboratory/splicing-pipelines-nf). Raw sequencing files generated in this study will be deposited on GEO upon manuscript acceptance.

